# Molecular Species Determination of Cyathostomins from Horses in Ireland

**DOI:** 10.1101/2023.12.13.571572

**Authors:** Orla Byrne, Disha Gangotia, John Crowley, Annetta Zintl, Liam Kiser, Olivia Boxall, Daniel McSweeney, Fiona O’Neill, Stacey Dunne, Breanna Rose Lamb, Nicola Walshe, Grace Mulcahy

## Abstract

Cyathostomins are globally important equine parasites, responsible for both chronic and acute pathogenic effects. The occurrence of mixed infections with numerous cyathostomin species hinders our understanding of parasite epidemiology, host-parasite dynamics and species pathogenicity. There have been few studies of cyathostomin species occurring in horses in Ireland, where temperate climatic conditions with year-round rainfall provide suitable conditions for infection of grazing animals with bursate nematodes. Here, we amplified and sequenced the ITS-2 region of adult worms harvested at post-mortem from 11 adult horses between August 2018 and June 2020, and recorded species prevalence and abundance of worms recovered from the caecum, right ventral colon and left dorsal colon, using both BLAST and IDTAXA for taxonomic attribution. Phylogenetic relationships and community composition were also recorded and compared with other relevant studies, including a global meta-analysis. Overall, our results agree with previous studies that there does not seem to be a major difference in cyathostomin species occurrence in equids in different geographical regions. We confirmed the results of other workers in relation to the difficulties in discriminating between *Cylicostephanus calicatus* and *Coronocyclus coronatus* on the basis of ITS-2 sequences.

## 1. Introduction

Cyathostomins are a group of parasitic helminths that inhabit the gastrointestinal tract. They are obligate parasites which are ubiquitous in equids and of significant pathogenic concern (Boxell *et al.*, 2004; Ghafar *et al.*, 2022; Hinney *et al.*, 2011; Love *et al.*, 1999; Morariu *et al.*, 2016; Steinbach *et al.*, 2006). Currently, there are over 50 recognised species with 40 recorded as infecting horses, and mixed infections are the norm (Lichtenfels *et al.*, 2008; Lyons *et al.*, 1999). Cyathostomins have a direct life cycle, with infection occurring through ingestion of infective stage (L3) larvae on pasture. Following infection, cyathostomin larvae penetrate into the mucosa of the large intestine, and can remain here in a dormant (hypobiotic) state, emerging to mature and establish patent infection in the lumen after a variable pre-patent period of up to three years (Gibson, 1953; Nielsen and Lyons, 2017). The hypobiotic state is relevant to anthelmintic efficacy and to cyathostomin-associated disease (Boisseau *et al.*, 2023; Morariu *et al.*, 2016). Typically, of the total worm burden in an individual animal, the majority will comprise hypobiotic larvae in the mucosa (Dowdall *et al.*, 2002; Duncan *et al.*, 1998; Gibson, 1953; Matthews, 2008). The stimuli responsible for hypobiotic larvae resuming development and completing their life-cycle are unclear, but are thought to involve cross-talk between adult and larval worms, along with interaction with the gut microbiota, the host immune system and environmental disruption such as anthelmintic treatment (Gibson, 1953; Love *et al.*, 1999; Matthews, 2008; Walshe *et al.*, 2019; Walshe *et al.*, 2021; Walshe *et al.*, 2020). The existence of a large number of species that are typically present in mixed infections highlights the importance of understanding these dynamics, in efforts both to understand the basic biology of cyathostomin infections and the provision of improved control strategies (Bellaw and Nielsen, 2020; Hodgkinson *et al.*, 2003).

Horses may tolerate light cyathostomin infection without the development of overt clinical signs (Love *et al.*, 1999; Morariu *et al.*, 2016; Murphy and Love, 1997; Walshe *et al.*, 2021). However, high burdens can lead to ill-thrift and poor performance (Murphy and Love, 1997) and in some cases a specific clinical presentation of acute larval cyathostominosis (ALC), which develops in response to synchronous emergence of larvae from the mucosa and involves severe local and systemic inflammation, with high morbidity and potential mortality (Hodgkinson *et al.*, 2003; Love *et al.*, 1999; Walshe *et al.*, 2020). The emergence of encysted cyathostomin larvae is believed to be affected by many factors, including negative inhibition from an established adult worm population, removed following anthelmintic treatment (Steinbach *et al.*, 2006; Walshe *et al.*, 2019; Walshe *et al.*, 2021). ALC has a 50% mortality rate recorded in referral centres (Giles *et al.*, 1985; Peregrine *et al.*, 2006; Walshe *et al.*, 2021).

In the past, characterisation of cyathostomin communities relied exclusively on morphological identification of species (Kinsella, 2002; Lichtenfels *et al.*, 2008). However, this is laborious, time-consuming and highly specialised (Avramenko *et al.*, 2015; Hodgkinson *et al.*, 2003; Lichtenfels *et al.*, 2008), and it has largely been supplanted by molecular species determination, either of individual worms, or using population analysis “Nemabiome” approaches (Avramenko *et al.*, 2015).

There are just two previously published studies on characterising cyathostomin species in horses in Ireland. The first (Kinsella, 2002), used morphological species determination and the second (Elghryani, 2018) used reverse line blots of the intergenic spacer gene region. Elsewhere, internal transcribed spacer 2 (ITS-2) primers as described by (Gasser *et al.*, 1993) have been widely used as taxonomic markers for cyathostomins. The focus of recent studies have utilised community analysis tools, such as ‘Nemabiome’ application of ITS-2 sequencing (Avramenko *et al.*, 2015; Courtot *et al.*, 2023; Halvarsson *et al.*, 2023; Nielsen *et al.*, 2022) .

We aimed to sequence the ITS-2 region from individual cyathostomins retrieved from naturally infected horses in Ireland in a post-mortem study and to compare the community composition from the colon and caecum to those described in previous studies in Ireland and internationally. We also aimed to explore any difficulties in distinguishing species based on ITS-2 sequencing, to investigate any trends in seasonality in the data, and to compare and contrast two different methods of inferring taxonomy based on ITS-2 sequences, using BLAST and IDTAXA methods.

## 2. Materials and Methods

### 2.1. Ethical approval

This was a post-mortem study and did not require any specific ethical approval.

### 2.2. Study population

We conducted an abattoir survey between August 2018 and June 2020. A total of eleven equine carcasses were selected at random for examination at two Irish abattoirs. In accordance with food safety legislation, no horse had received anthelmintic treatment within 32 days of slaughter. The horses were assessed for clinical abnormalities by a veterinarian prior to inclusion in the study as part of the normal pre-mortem inspection at the abattoir.

### 2.3. Sample processing

Each carcass was processed as per abattoir standard operating procedures. Sampling was undertaken on the same day as euthanasia. To isolate the large intestine, the ileocecal junction was located, tied off with cable ties and dissected. The liver, stomach, spleen, pancreas, and small intestine were removed, and the large intestine was transferred from the processing line to a dissection table.

Further cable ties were placed at the junction between the caecum and right ventral colon, the junction between the right ventral and left ventral colon, the junction between the left ventral colon and left dorsal colon, and the junction between the left dorsal and right dorsal colon. An incision was made in each of the three intestinal segments (caecum, right ventral colon (RVC) and left dorsal colon (LDC), and the luminal contents were emptied separately into individual 10L graduated buckets. For each section, the mucosa was gently rinsed, and the liquid was added to the bucket containing luminal contents. The contents were then stirred whilst water was added to make a homogeneous suspension to a volume of 10L. Two 100ml aliquots were taken from each section, equating to 2% of the total volume. Faeces were collected from the rectum of each horse and placed in a ziplock bag. All samples were placed on ice for transfer to the laboratory.

### 2.4. Faecal Egg Counting

The Mini-FLOTAC technique (Cringoli, 2006) was utilised to count strongyle eggs in faeces. Briefly, 45ml of saturated NaCL was added to 5g of faeces and the flotation chambers were completely filled with the faecal suspension. After 10 minutes, the key was used to turn the reading disc clockwise (about 90°), the sample was examined under the microscope and the eggs per gram calculated. The sensitivity limit for the Mini-FLOTAC is 5 epg.

### 2.5. Luminal worm retrieval

Each aliquot representing 2% of the contents of the caecum, RVC and LDC was examined individually using a modification of the Baermann method by enclosing them in a muslin parcel suspended in water in a conical flask and leaving to sediment for 24 hours. The entire sediment was then examined in detail (using aliquots forming a thin layer in a petri dish) under a dissecting microscope. The remaining sediment was also examined under a dissecting microscope.

The total number of worms present in each 200ml sample was enumerated and recorded. Individual worms were removed manually and washed prior to storage. Each worm was placed in a 2ml Eppendorf with >/=50ml PBS and stored at -20°C. For each section a maximum of 50 worms (range 1-50) was retained for subsequent examination from combined duplicate samples. IBM SPSS (Version 27) was used for descriptive statistics. The Kruskal-Wallis test was utilised for non-parametric data including faecal egg counts (FEC), numbers of worms and species distribution

### 2.6. DNA Extraction and amplification

#### 2.6.1. Genomic DNA extraction from individual nematodes

Individual worms were dissected into a minimum of three parts using a sterile needle. DNA was then extracted from each worm using the Powerfecal Pro extraction kit (Qiagen, Product number 51804), as per the standard operating protocol, with minor adaptations. Specifically, samples were pulse vortexed, as part of the bead beating step, for a minimum of 45 minutes, the dry spin after solution C5 was at 16,600 x G for three minutes, and solution C6 of the kit was put through the filter membrane twice to increase elution yield (Bredtmann *et al.*, 2019). After extraction, samples were stored at -20°C until PCR amplification.

#### 2.6.2. PCR amplification

PCR reactions were performed in 50μl reaction volumes using ITS-2 primers as described by Gasser *et al.*, (1993), (NC1 forward; 5’ACGTCTGGTTCAGGGTTGTT3’ and NC2 reverse; 5’TTAGTTTCTTTTCCTCCGCT3’).

The reaction mixture consisted of 10μl of Promega GoTaq Flexi Buffer,(product no. M780A) 1.5mM of Promega MgCL2 solution, 0.2mM Promega DNTP PCR nucleotide mix, (product number U151A) 1μM of each forward and reverse primer (Eurofins Genomics) 1.25U of Promega GoTaq G2 Flexi DNA Polymerase, (product no. M780A) 10μl template DNA and 15.75μl nuclease free water (Merck Millipore 693520). Amplification steps included initial denaturation at 95°C for 2 minutes, then cycling steps starting at 95°C for 1 minute, annealing at 55°C for 1 minute and extension at 72°C for 1 minute, for 40 cycles, followed by extension at 72°C for 5 minutes.

For each run, a negative control containing 10μl nuclease-free water in place of DNA template was included.

Amplicons were purified using the Qiagen Qiaquick PCR purification kit (Product.No. 28106) and analysed through Sanger sequencing in both directions using the PCR primers. Sequencing was outsourced to Eurofins Genomics, (Sandyford, Dublin 18, Ireland) and FASTA files were returned to the research group for analysis.

### 2.7. Molecular taxonomy attribution

Aligned sequences were constructed from forward and reverse reads and compared with published nematode sequences in the NCBI database using Clustal Omega.

Samples that did not yield a reliable BLAST result, (E value > E^-10^, low query cover and percentage identity <95%,) were removed from the dataset.

As we did not carry out morphological examination of recovered worms, all sequences are uploaded as ‘unidentified cyathostomin’ on NCBI database and a note accompanying each species outlining its closest hit on BLAST. These nucleotide sequence data is available in the GenBank™ database under the accession numbers: **<u>OR941129 - OR941242.</u>**

#### 2.7.1 Analysis of sequences using Nematode ITS2 database and IDTAXA

Sequences obtained from individual worms were also compared with data in an opensource ITS2 rDNA database for both parasitic and free living nematodes (Wokentine *et al.*, 2020). To infer taxonomy, IDTAXA (Murali *et al.*, 2018) as a component of the DECIPHER (2.22.0) package in R (v4.1.2), and phyloseq package version (1.38.0) through R studio (v2023.12.0.369) was used within the DADA 2 pipeline as described at www.nemabiome.ca.

### 2.8 Phylogenetic analiysis

MEGA11, and iTOL (https://itol.embl.de/) were used for phylogenetic tree creation. MEGA11 was used for MUSCLE alignment and trimming of sequences. A tree was created using the Maximum likelihood algorithm with 1,000 bootstrap replications and pairwise deletion for gaps. (Tamura *et al.*, 2021). The evolutionary history was inferred by using the Maximum Likelihood method and Tamura-Nei model(Tamura and Nei, 1993). The bootstrap consensus tree inferred from 1000 replicates (Felsenstein, 1985) is taken to represent the evolutionary history of the taxa analyzed. Branches corresponding to partitions reproduced in less than 50% bootstrap replicates are collapsed. Initial tree(s) for the heuristic search were obtained automatically by applying Neighbor-Join and BioNJ algorithms to a matrix of pairwise distances estimated using the Tamura-Nei model, and then selecting the topology with superior log likelihood value. Seven sequences were removed at the alignment section of the phylogenetic analysis as they were too short when aligned with other species. There were a total of 203 positions in the final dataset. Visualisation of the phylogenetic tree was conducted using iTOL V6.8.1(Letunic and Bork, 2021)

## 3. Results

### 3.1. Study cohort and worm recovery

Table 1 provides the details of horses included in the study, together with their strongyle FEC. number of worms, and the areas from which worms for molecular identification were retained. The horses included in the study ranged from yearlings to twenty-three years old with a mean of 7.7 years of age, and a median of 6 years. There were four females and seven males. Five were Thoroughbred, and six Irish Sport Horse. FEC ranged from 0-1,670 eggs per gram (epg). Nematodes were retrieved from two horses with a FEC of 0. Conversely, high FEC did not always correlate with high luminal burden of nematodes. Numbers of worms recovered from the caecum were lower than those recovered from LDC (P=0.009). No other significant differences in worm recovery between compartments was noted. Up to 14 individual worms from each intestinal compartment, per horse, had taxonomy inferred to species level by BLAST and IDTAXA. For all but one horse, worms from two or three compartments were included in the taxonomic analysis.

**Table 1:** Horse details, FEC and number of worms recovered and speciated from each intestinal compartment. RVC= Right Ventral Colon and LDC = Left Dorsal Colon. * = P < 0.05. Mean calculations are +/- the SEM (standard error of the mean). The number of worms recovered accounts for a 1% subsection of the total luminal contents of each animal sampled. Numbers of worms recovered from the caecum were lower than those recovered from LDC (P=0.009). No significant difference was recorded between RVC or from composite RVC and LDC colon to the caecum (P=0.291). FEC represents eggs per gram (epg) of faecal material).

| Horse | Horse Details |  |  |  |  | Number of worms recovered |  |  | Speciated Worms |  |  |
| --- | --- | --- | --- | --- | --- | --- | --- | --- | --- | --- | --- |
|  | Age | Breed | Sex | Sampling Date | FEC (epg) | RVC | LDC | Caecum | RVC | LDC | Caecum |
| 1 | 16 | ISH | F | Sep-18 | 1550 | 79 | 188 | 10 | 2 | 9 | - |
| 2 | 8 | ISH | M (gelding) | Sep-18 | 0 | 11 | 3 | 1 | 5 | - | - |
| 3 | 1 | TB | M (colt) | Sep-18 | 1670 | 205 | 155 | 12 | 8 | 2 | 4 |
| 4 | 10 | ISH | F | Oct-18 | 25 | 201 | 245 | 23 | 3 | 13 | - |
| 5 | 1 | TB | M (colt) | Oct-18 | 905 | 590 | 33 | 16 | 3 | - | 4 |
| 6 | 6 | ISH | M (colt) | Feb-20 | 225 | 2 | 26 | 0 | 1 | 4 | - |
| 7 | 2 | TB | M (colt) | Feb-20 | 800 | 1 | 249 | 2 | - | 4 | 1 |
| 8 | 12 | ISH | M | Mar-20 | 50 | 0 | 2 | 0 | - | 2 | - |
| 9 | 23 | TB | F | Mar-20 | 85 | 9 | 24 | 1 | 3 | 3 | - |
| 10 | 3 | TB | F | Jun-20 | 715 | 1410 | 3328 | 33 | 8 | 5 | 14 |
| 11 | 3 | ISH | M | Jun-20 | 0 | 40 | 56 | 53 | 4 | 5 | 9 |
| Total | - | - | - | - | - | 2548 | 4309 | 151 | 37 | 47 | 32 |
| Mean | 7.7+/-<br>2.12<br>4 | - | - | - | 547.73 +/-<br>188.83 | 231.63 +/-<br>129.21 | 391 +/-<br>295* | 13.73 +/-<br>5.1* | 3.36 +/-<br>0.83 | 4.27 +/-<br>1.16 | 2.91 +/-<br>1.4 |
| Range | 1-23 | - | - | - | 0-1670 | 0-1410 | 2-3328 | 0-53 | 0-8 | 0-13 | 0-14 |

### 3.2. BLAST analysis of ITS-2 amplified sequences

A total of 116 samples were extracted, amplified and compared using the BLAST algorithm with sequences in the NCBI database. Another 43 samples did not provide adequate sequence information for analysis Of the 116 sequences analysed, 43 samples had a 100% identity score with their presumed species. The remaining samples had taxonomy inferred based on similarity scores of > 95% similarity, and E,= 10 ^-10^ . The amount of samples that fell within the percentage similarity values outlined previously (Mitchell *et al.*, 2019) are outlined as follows: >99% n= 46, >98% n= 16, >97% n= 5, >96% n= 3.. (Supplementary File).

### 3.3 IDTAXA analysis of amplified sequences and comparison with BLAST results

A total of 103 of the original 116 consensus Sanger sequences gave valid outputs from the nemabiome.ca workflow using the ITS2 database using IDTAXA to infer taxonomy. Thirteen sequences did not pass quality control cut-offs for the IDTAXA algorithm because they fell below the 60% taxonomic classification confidence threshold. Of the 103 sequences meeting the threshold, 83 (80.5%) were assigned to species level by IDTAXA and gave the same result as BLAST using >95% similarity, (Table 2). Of the remainder, 13 were identified only to genus level by IDTAXA, but as the same genus as BLAST. A further six were presumptively identified as *Cylicostephanus calicatus* by BLAST and *Coronocylus coronatus* by IDTAXA. One sequence identified on BLAST as *Cor. coronatus* was assigned as unclassified *Cyathostomum* spp. using IDTAXA.

**Table 2.** Comparison of BLAST and IDTAXA identification of ITS sequences. We compared the degree to which BLAST, with >95% sequence identity, and the IDTAXA algorithm gave comparable results using our ITS-2 sequences.

| Sequences for which taxonomy inferred by BLAST | Agreement on Species using IDTAXA | Agreement on Genus level, IDTAXA | Inability to distinguish between *Cor. coronatus* and *Cyl. calicatus* | Species BLAST, different genus IDTAXA | Did not pass IDTAXA Quality Threshold |
| --- | --- | --- | --- | --- | --- |
| ***Cor. coronatus*** | 26 | 2 | 7 | 1 (unclassified Cyathostomum). | 5 |
| ***Cyl. nassatus*** | 15 |  |  |  |  |
| ***Cyl. goldi*** | 11 |  |  |  |  |
| ***Cyl. insigne*** | 11 |  |  |  |  |
| ***Cyl. longibursatus*** | 9 | 1 |  | 3 | 3 |
| ***Cyl. ashworthi*** | 3 | 6 |  |  |  |
| ***Cya. pateratum*** | 2 | 1 |  |  |  |
| ***Cya. catinatum*** | 2 | 3 |  | 2 | 2 |
| ***Cor. labiatus*** | 1 |  |  |  |  |
| ***Cyl. leptostomus*** | 1 |  |  |  |  |
| ***Cyl. minutus*** | 1 |  |  |  |  |
| ***S. vulgaris*** | 1 |  |  |  |  |
| ***Cyl. calicatus*** |  |  |  |  | 2 |

### 3.4. Species recovered, abundance and prevalence

The species abundance in each of the three intestinal compartments is shown in Figure 1. *Cor. coronatus* was the most abundant species recovered (29%). Between them, *Cor. coronatus*, *Cylicocyclus nassatus* and *Cylicostephanus longibursatus* accounted for more than half (54%) of the entire dataset. Although there was no significant difference in species distribution across the three intestinal compartments, *Cylicocyclus insigne* was equally abundant to *Cor. coronatus* in the LDC. This compartment also had the most diverse range of species (n=11), with the caecum having the least diversity (n=8).

**Figure 1:**
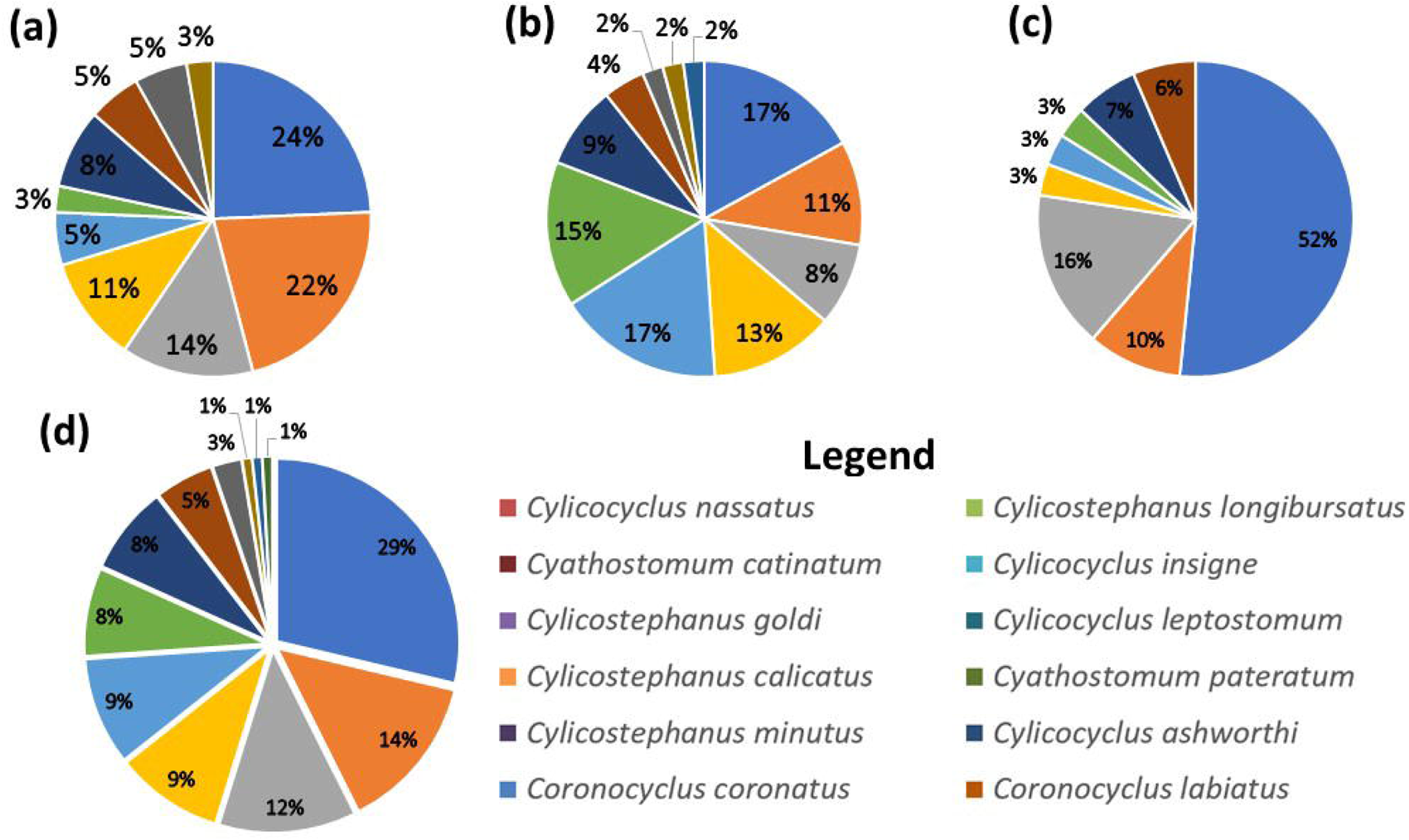
Species abundance data from samples obtained from Right Ventral Colon (a) Left Dorsal Colon (b), Caecum (c) and across all three compartments (d). *Cor. coronatus* was the most abundant species recovered, at 29% of total nematodes speciated. Between them, *Cor. coronatus, Cyc. nassatus* and *Cyc. longibursatus* accounted for more than half (54%) of the entire dataset. The LDC had the most diverse range of species (n=11), with the caecum having the least diversity (n=8).

Figure 2 outlines the composition per horse of presumed species. It outlines the species found in each of the 11 horses, represented as a percentage of the total samples amplified from each individual. The actual number of worms that each segment represents is also included in the figure, overlaying each segment. This figure provides an insight into the species composition differences between individuals from the samples that were obtained in this study. For the purposes of this analysis, *Strongylus vulgaris* (horse 5) was included in this analysis.

**Figure 2:**
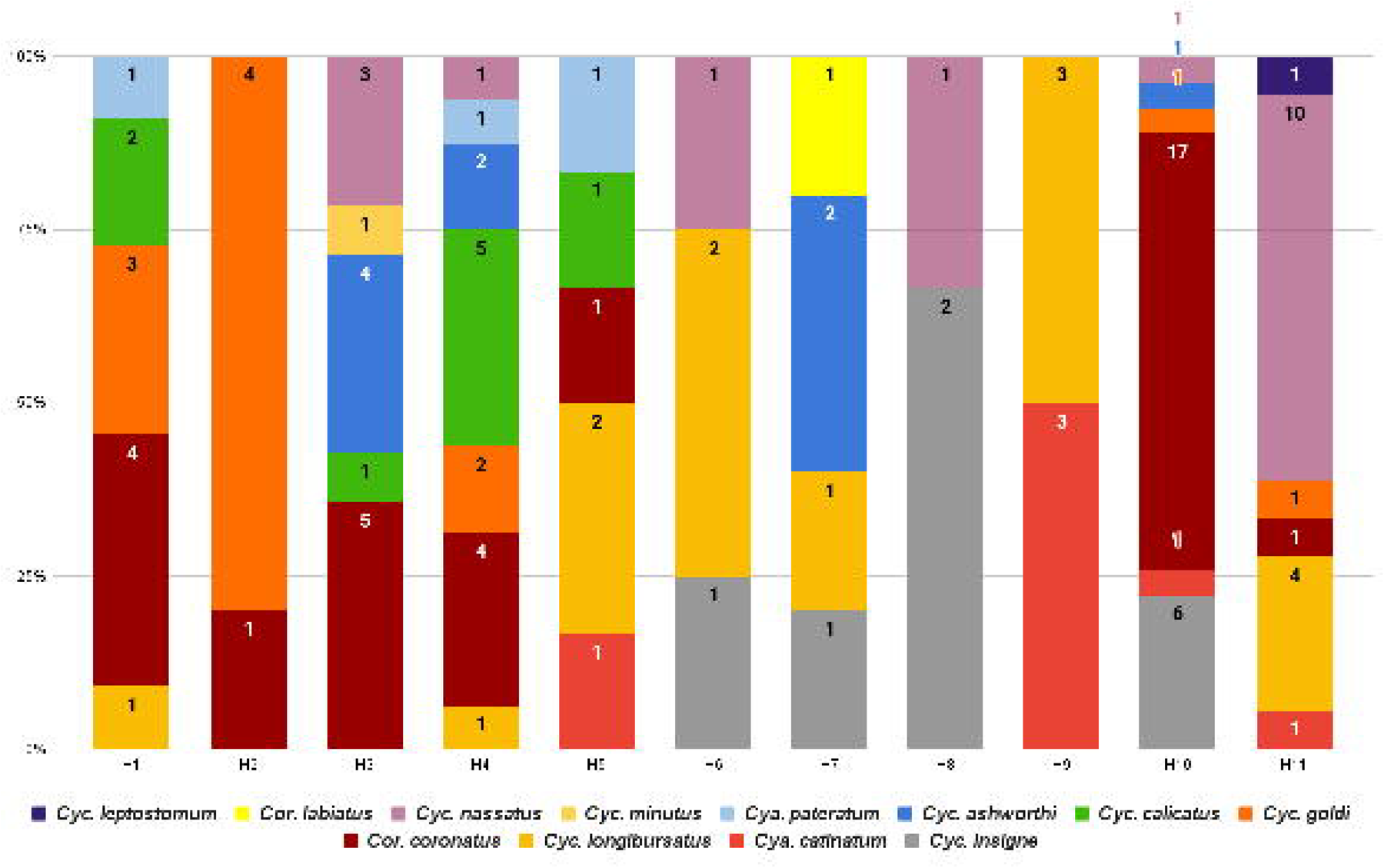
100% stacked bar graph of the species distribution per horse. Species richness ranged from 2-7 per horse. Each horse represents a stacked column, identified on the X-axis. The species are represented by coloured sections illustrated as a percentage of the nematodes obtained from each horse. The actual number of samples from each species is also illustrated within each segment.

The prevalence of individual species and the richness of cyathostomin communities in this study population is illustrated in Figure 3. *Cor. coronatus* and *Cyc. longibursatus* were the most prevalent, found in seven of the eleven horses (63.63% prevalence), *Cyc. nassatus* was present in six horses (54.54%) and *Cylicostephanus goldi* in five horses (45.45%). *Cyc. insigne, Cylicostephanus calicatus, Cylicocyclus ashworthi* and *Cyathostomum catinatum* were each recorded in four horses, at a prevalence of 36.36%. *Cyathostomum pateratum* was found in three horses (27.27% prevalence) with *Cylicostephanus minutus, Coronocyclus. labiatus* and *Cylicocyclus leptostomum* in one animal each, comprising 9.09% prevalence.

**Figure 3:**
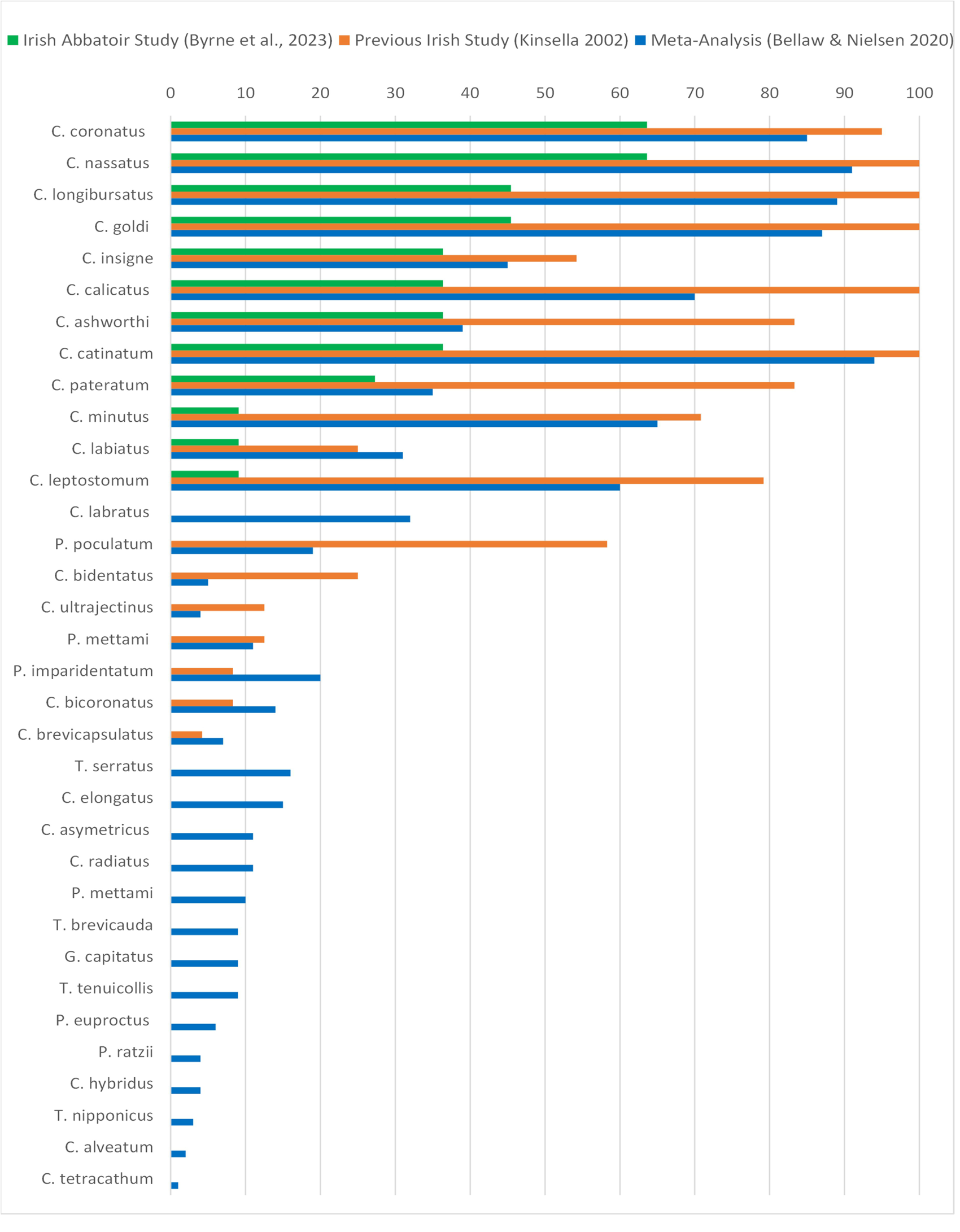
Prevalence of cyathostomin species as compared with a previous study in Ireland (Kinsella 2002) and a global meta-analysis (Bellaw and Nielsen, 2020). We identified sequences similar to twelve cyathostomin species recorded in both the previous Irish study and the global meta-analysis. There were no species identified in either study in Ireland that were not identified within the global meta-analysis. Generally, the most prevalent species in the two previous studies were also highly prevalent in our study.

Minimal richness from the samples taken ranged from two to seven species per individual, with a mean of 4.27 +/-0.604 (SEM). The modal species richness was two and five species equally, with a median species richness of five within this dataset. Over 50% (54.5%) of the horses sampled had >5 cyathostomin species present. One large strongyle, *Strongylus vulgaris,* was identified (Horse 5) but not included in the dataset.

### 3.5 Analysis in relation to previous local and global data

Next, (Figure 4) we compared the composition of cyathostomin species found in our study with data from a previous study in Ireland (Kinsella, 2002) and a global meta-analysis (Bellaw and Nielsen, 2020). Twelve cyathostomin species were common to our study, Kinsella, 2002, and the global meta-analysis. Notably, *Cor. labratus* was not identified in either of the Irish datasets, but features in the global meta-analysis. Our study presumptively identified 12 of the 13 most prevalent global species, (Bellaw and Nielsen, 2020), using two different methods of inferring taxonomy from ITS-2 sequences.

**Figure 4:**
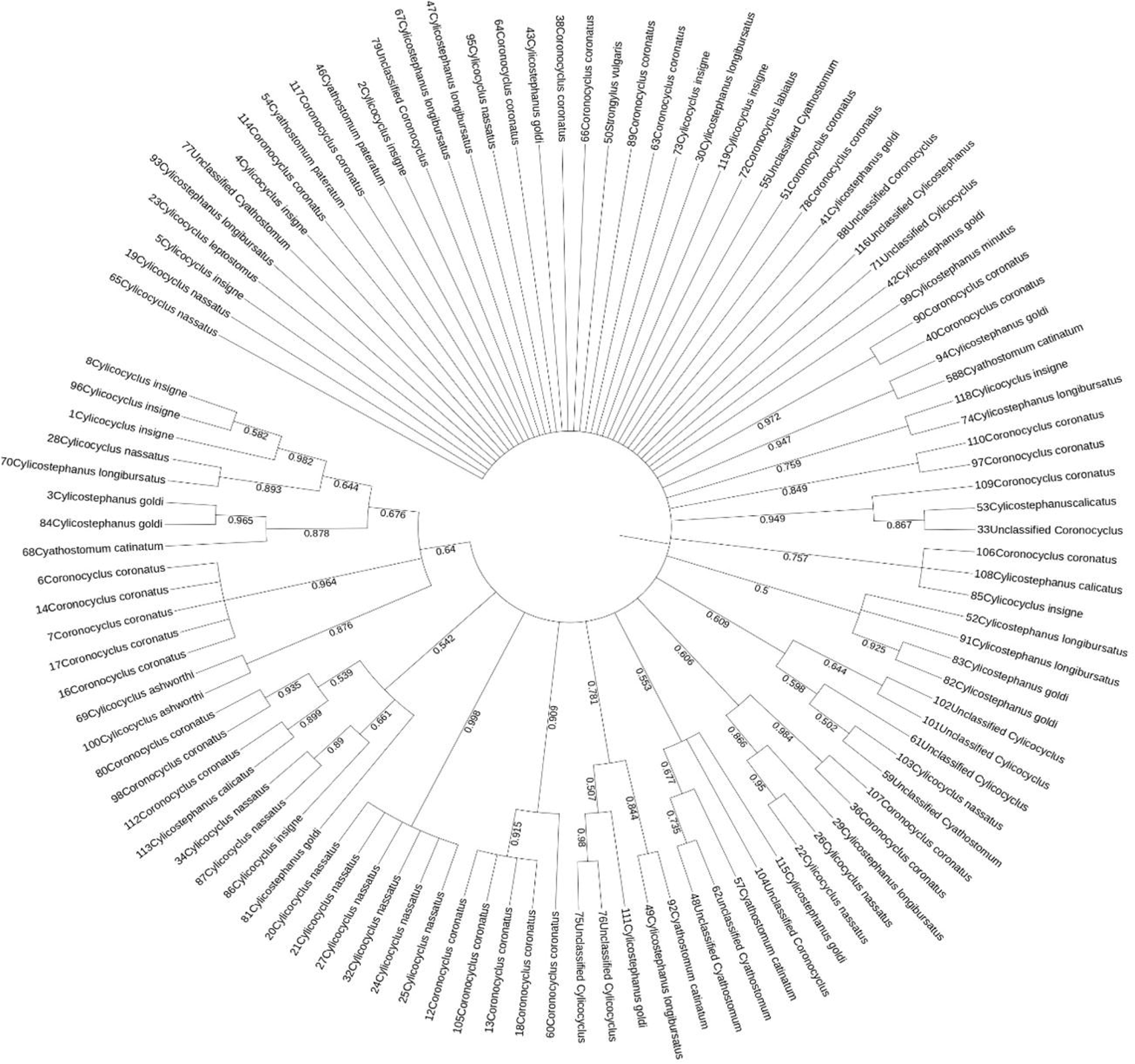
Full phylogenetic tree of sequences from recovered worms. IDtaxa similarity sequences for individual cyathostomins were used for phylogenetic analysis. MEGA11 was used for MUSCLE alignment and trimming of sequences. A tree was created using the Maximum likelihood algorithm with 1,000 bootstrap replications and pairwise deletion for gaps. This tree unveiled species that located within groupings of other species, notably *Cylicostephanus calicatus* and *Coronocyclus coronatus*.

### 3.6. Phylogenetic analysis of species

Sequences were used for construction of a Maximum Likelihood phylogenetic tree (Figure 5) which was rooted at the midpoint. Figure 5 unveiled some interspecies occurrences within branches, notably between *Cyc. calicatus* and *Cor. coronatus*.

## 4. Discussion

There was no direct relationship between strongyle faecal egg count and worm burden in the carcases examined, which was not unexpected (Chapman *et al.*, 2003a; Chapman *et al.*, 2003b; Nielsen *et al.*, 2010). We recovered worms from horses with a 0 EPG count, which has been well-documented previously (Gibson, 1953; Love *et al.*, 1999; Matthews, 2008; Walshe *et al.*, 2019; Walshe *et al.*, 2020).

We did not attempt morphological identification of the worms recovered in our study, because of the inherent difficulties in doing so and our lack of expertise in this area. Our taxonomic classifications, therefore, are inferred solely on the basis of sequence information. Clearly, reliance on molecular data only for this purpose raises the possibility of errors due to errors in sequence databases, judgement calls when there is similarity to more than one species, and the inability to distinguish between some species on the basis of one locus (in this case ITS-2) alone. In order to validate and provide for greater reliability from our results, we used two distinct methods for inferring taxonomy from sequence information, BLAST, and IDTAXA.

IDTAXA is a method of taxonomic classification based on sequences of marker genes from organisms found in microbiome samples or other sources. It involves assigning sequences to a reference taxonomy containing sequences from known members of a reference dataset, and avoids overclassification of sequences from organisms which may not be present in the reference set (Murali *et al.*, 2018). A database of ITS2 rDNA sequences from both parasitic and free-living nematodes is publicly available, (Workentine *et al.*, 2020) and is now widely used in “nemabiome” studies in order to characterise a mixed population of parasitic nematodes within a single host, or host population, including in horses (Ghafar *et al.*, 2022; Halvarsson *et al.*, 2023; Poissant *et al.*, 2021; Sargison *et al.*, 2022) following high-throughput parallel sequencing and metabarcoding. However, the same bioinformatics pipeline using DADA 2 can be used to assign taxonomy to data obtained by conventional Sanger sequencing. The results obtained in our study from BLAST of individual sequences and from the DADA 2 IDTAXA output gave identical results to species level for 82 (80.6%) of worms and results consistent to genus level for a further 12 (12.6%). For the remaining seven sequences, (6.8%) there was inconsistency in distinguishing between *Cor. coronatus* and *Cyl. calicatus*, reflecting difficulties in using ITS-2 sequences to distinguish between these potentially cryptic species in other studies (Bredtmann *et al.*, 2019,(Louro *et al.*, 2021).

Overall, given the consistency between taxonomic assignment by BLAST, and by IDTAXA, in this study, together with the difficulty and potential for error in morphological identification of strongyle worms of horses, we suggest that comparison of two sequence-based methods of taxonomic classification is useful in further determining the taxonomic relationships between these parasites. The characterisation of additional loci may also be useful in some cases (Courtot *et al.*, 2023)

This study detected sequences similar to twelve of the thirteen most common species globally (Figure 4), as compared to Bellaw and Nielsen’s meta-analysis (Bellaw and Nielsen, 2020). It should be noted that prevalence as outlined in this study is based on a 2% sample of contents of each intestinal compartment sampled and thus is likely to underestimate prevalence. Among the nineteen species identified in the morphological study by Kinsella (2002), we detected twelve (Figure 4). Notably, the prevalence of *Cya. catinatum* was 36% in our study, in contrast to 100% prevalence in Kinsella’s work and 94% in the global meta-analysis. Elghryani (2018), found *Cyc. nassatus* as the most prevalent among the twelve species identified. The high abundance and prevalence of *Cyc. nassatus,* occurring in 64% (n=7) of the horses sampled, is consistent among all three Irish studies and the global meta-analysis. The prevalence findings between the three studies from Ireland broadly reflect one another. A notable observation was the absence of the twelfth most prevalent species as identified by (Bellaw and Nielsen, 2020), *Cor. labratus,* which was absent from the Irish studies, and has not been detected either in some studies in similar climates-Wales and Scotland(Mitchell *et al.*, 2019; Sargison *et al.*, 2022). This species has however, been recorded in England and France (Cwiklinski *et al.*, 2012; Peachey *et al.*, 2018).

*Cyc. calicatus* and *Cor. coronatus* are interspersed within the ML tree. This reflects their close relationship, as previously identified, and the limitations of using the ITS-2 locus to differentiate the subsets of these species (Courtot *et al.*, 2023; Hung *et al.*, 2000; Louro *et al.*, 2021).

Across the three intestinal compartments examined in this study, the number of worms retrieved from the caecum was significantly lower than that recovered from LDC (P =0.009). This was also observed in a previous Irish study(Kinsella, 2002). (Ogbourne, 1976), assessed abundance of cyathostomins postmortem and found 50% in the ventral colon and 45% in the dorsal colon. The worm burden in the caecum represented only 5% of the overall cyathostomin burden. Similarly, (Stancampiano *et al.*, 2010), observed that the majority of small strongyles were concentrated in the ventral colon (67.34%) and less frequently in the caecum (27.52%) and dorsal colon (5.14%). While this distribution was also reflected by Kinsella (2002), the converse was found for encysted stages of cyathostomins, as recorded by (Nielsen *et al.*, 2021) who documented 40% of EL3/LL3/L4 in the caecum, and 38%/22% in the ventral and dorsal colon respectively. This recurrent recording of biological preference of luminal cyathostomins for the colon over the caecum, and the inverse for encysted larvae may reflect the slower transit time of the colon and the natural peristalsis of food through the digestive tract, enabling passive transport and a longer exposure to the intestinal wall necessary to facilitate larval encystment (Geor *et al.*, 2013; Miyaji *et al.*, 2008).

Over half of the horses in this study had more than five cyathostomin species recovered from them (Figure 2).(Abbas *et al.*, 2023; Reid *et al.*, 1995)

## 5. Limitations

The most important limitation of this study is sample size. This is a descriptive study, which affects the potential for statistical power. Conclusions from observations within any study with a limited sample size must take this into account. Results are considered in the context that only two nematodes were retrieved for sampling from horse eight. Notably, it is clear from this research and other studies that the ITS-2 locus cannot resolve all isolates of *Cor. coronatus* and *Cyc. calicatus,* pointing to the need for a different locus for further molecular work, such as COX1, or a multi-locus approach (Courtot *et al.*, 2023; Ghafar *et al.*, 2022). We acknowledge also the value of examining seasonality in relation to cyathostomin species prevalence and abundance, as well as potential interactions, positive or negative between species. Variability of seasonal patterns among species, in particular, is an understudied facet of cyathostomin biology which may be crucial for comprehensive characterisation of pathogenesis and epidemiology of cyathostomin-related disease, as this tends to have a seasonal distribution of presentation(Abbas *et al.*, 2023; Reid *et al.*, 1995). Other studies, (Salle *et al.*, 2018) have begun to explore interactions among cyathostomin communities, and this also has a potential bearing on the pathogenesis and epidemiology of cyothostomin-related disease. Our study was too small to make reliable observations on these points.

## 6. Conclusions

This study has produced new insights into cyathostomin species occurrence, interactions and, community composition in horses in Ireland, with comparisons to previous studies in this locality and a global meta-analysis. It also displays a high level of agreement in assigning taxonomy from ITS-2 sequences, based on BLAST and IDTAXA, which we believe is useful in increasing the reliability of studies using molecular information to study cyathostomin communities.

Post-mortem analysis and identification in naturally-infected hosts, as conducted here, will continue to have a role to play in advancing understanding of cyathostomin biology and pathogenesis, in parallel with population-based approaches. More extensive studies involving both population and post-mortem studies are warranted in order to enhance risk assessment, biomarker-based diagnostic tools and treatment protocols (Walshe *et al.*, 2020). It is also useful in investigating potentially cryptic species as seen in this study with *Cor. coronatus* and *Cyc. calicatus*.

Studies on cyathostomin population dynamics are a crucial step in understanding the complex picture of cyathostomin infections in the face of pervasive anthelmintic resistance and their potential to advance mechanistic understanding of gut health in horses.

## Author contributions

Conceptualization, G.M., N.W., O.B.; writing-original draft preparation, O.B. – review and editing, G.M., N.W., O.B. and A.Z.; laboratory work and bioinformatics-O.B., D.G., J.C.; laboratory contributions, L.K., O.BX., D.McS., F.ON., S.D., BR.L. All authors have read and agreed to the published version of this manuscript.

## Acknowledgements

We would like to acknowledge the valuable input and assistance of Amanda Lawlor, Senior Technical Officer, and Dr. Nagwa Elghyrani, Adjunct Research Fellow, UCD School of Veterinary Medicine

---

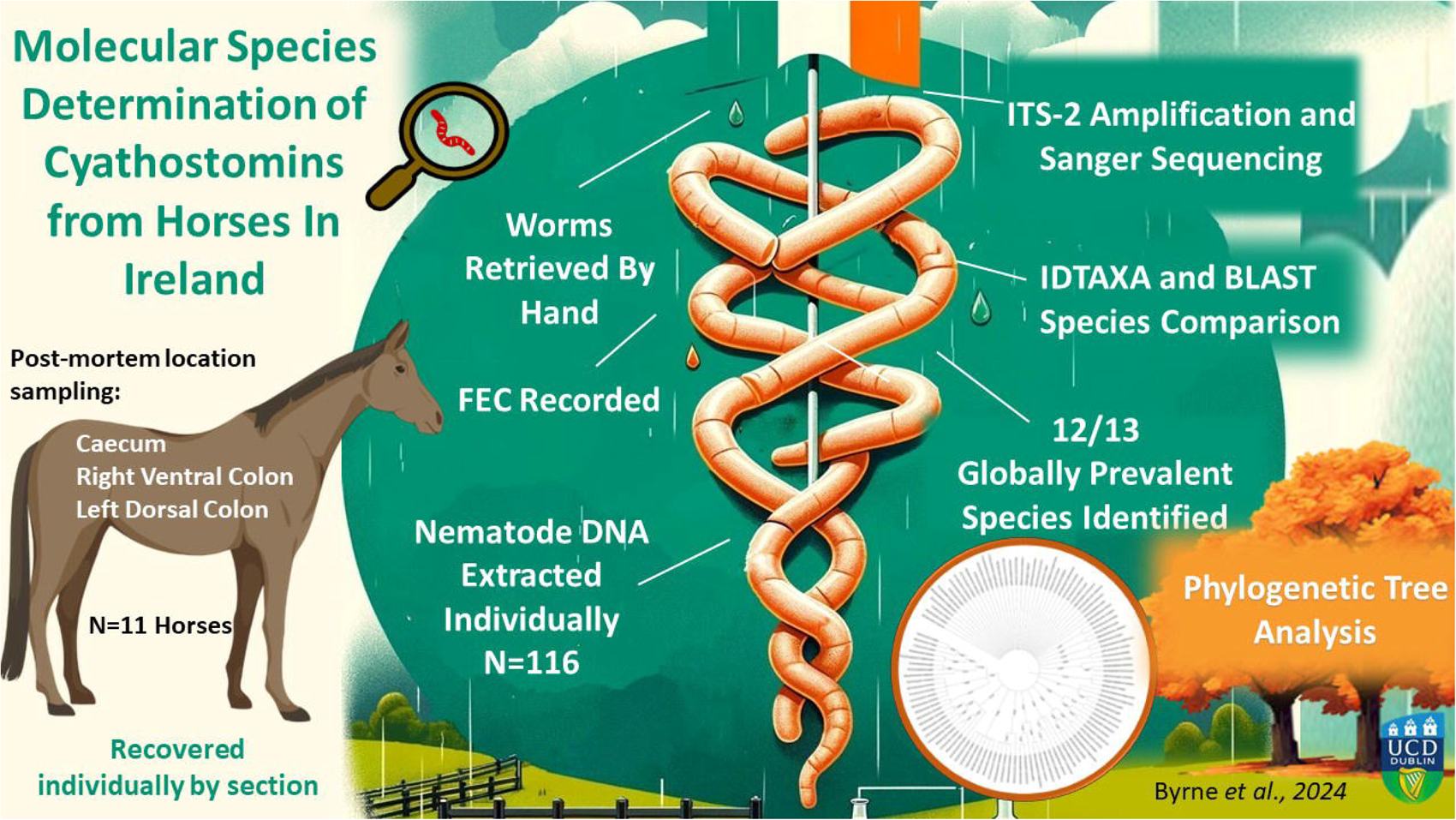

